# Medical diagnostic radiation promotes murine Apc/Kras-driven colon carcinogenesis

**DOI:** 10.64898/2026.06.11.731293

**Authors:** Mélissandre Gomot, Nora Essakhi, Clément Codan, Tasneem Abaza, Meryem Ulas, Marie Thérage, Chloé Brizais, Florence Bachelot, Amandine Sache, Karen Chaaya, Imène Garali, Quentin Pascal, Dmitry Klokov, Guillaume Varès

## Abstract

Computed tomography (CT) is one of the most widely used diagnostic imaging modalities worldwide, yet the biological risks associated with such exposures remain incompletely understood. Here, we investigated the effects of clinically relevant low (25 mGy) and moderate (250 mGy) dose radiation exposures on colon carcinogenesis using *in vivo* KPC:APC transgenic mice and an *ex vivo* organoid system carrying inducible *Apc* and *Kras* driver mutations. While medical diagnostic radiation did not initiate carcinogenesis in wild-type and did not alter carcinogenesis in *Apc*-mutant tissues, it significantly promoted the progression of precancerous lesions in the presence of both mutations, especially when exposure occurred during early tumor initiation. Organoids derived from mice harboring both *Apc* and *Kras* mutations mirrored this susceptibility, exhibiting radiation-induced enlargement and activation of transcriptomic and proteomic programs associated with colorectal cancer, including cell-cycle dysregulation and mTORC1 pathway activation. These findings show that radiation doses within the range delivered by routine abdominal CT imaging can potentiate the carcinogenic processes in genetically predisposed cells, underscoring the need to consider individual susceptibility when evaluating the benefit–risk balance of medical diagnostic exposures.

## 1. Introduction

Colon cancer (CC) is the third most commonly diagnosed cancer worldwide. In 2022, an estimated 1.1 million new cases were reported, resulting in more than 500,000 deaths globally [1]. Its development can be influenced by multiple factors. Those include non-modifiable determinants such as individual characteristics, medical history or genetic factors, as well as lifestyle-related factors (diet, smoking status, sedentary lifestyle, etc.) [2]. Among risk factors, ionizing radiation (IR) is infrequently highlighted, despite its increasing relevance in modern clinical practice. Over the last two decades, the global cumulative exposure to IR has doubled, largely driven by the use of computed tomography (CT) [3].

Approximately 400 million CT examinations are performed worldwide each year [4]. This corresponds to an average of 115.83 CT scans by 1,000 individuals, with substantially higher frequencies observed in high-income countries. According to the 2022 United Nations Scientific Committee on the Effects of Atomic Radiation (UNSCEAR) survey on diagnostic radiology, 52.24% of reported CT procedures included the abdomino-pelvic region in the targeted area, thereby exposing the colon to IR [5]. Dose levels varied considerably depending on the procedure type, scanner model, acquisition parameters, institutional protocols, and patient characteristics [6]. Mean effective doses among surveyed countries were 10.3 mSv for abdomen CT (liver, pancreas, kidneys) and 16.7 mSv for trunk CT (chest, abdomen, pelvis), with significant variability, reaching up to 30 mSv [5]. A subset of patients undergo multiple repeated examinations and accumulate cumulative effective doses exceeding 100 mSv [7].

Estimations of lifetime cancer risk from CT scans in the US suggested that annual CT use in 2023 might result in 103,000 future cancers, with lung and colon cancers the most common projected cancer types [8]. Recent epidemiological studies generally conclude that despite the estimated risk of solid cancer, diagnostic benefits outweigh potential adverse effects [9]. However, the influence of low-dose radiation (LDR) on pre-existing or early-stage malignancies remains poorly understood. Experimental investigations into the effects of low-to-moderate X-ray doses on colon carcinogenesis remain limited and have produced inconclusive findings regarding tumor initiation and progression [10,11]. At the cellular level, LDR may enhance proliferation and stem-like characteristics [12,13] yet the underlying mechanisms remain largely unresolved.

Here, we evaluated the specific effects of clinically relevant low dose medical diagnostic radiation on colon carcinogenesis using KPC:APC transgenic mice and a novel KPC:APC organoid model that harbor two key CC driver mutations, *Apc* and *Kras*. We characterized LDR-induced histopathological effects and explored the associated molecular mechanisms through transcriptomic and proteomic analyses. Additionally, we examined the role of N6-methylaadenosine (m6A) RNA methylation in the post-transcriptional regulation of radiation-modulated carcinogenesis.

## 2. Material and methods

### Animal experiments

Trios of B6.Cg-*Kras^tm4Tyj^ Apc^tm1Tno^* Tg(CDX2-cre/ERT2)752Erf/MaraJ (referred as KPC:APC) were purchased from the Jackson Laboratory (Bar Harbor, ME, USA). A colony was established and maintained in the ASNR pathogen-free animal facility by breeding Apc^fl/fl^ Kras^fl/+^ Cre^+^ mice with Apc^fl/fl^ Kras^+/+^ mice and Apc^fl/fl^ Kras^+/+^ Cre^+^ mice with Apc^fl/fl^ Kras^fl/+^ mice. The animals were housed in groups of 2 to 5 in individually ventilated cages with ad libitum access to food and water in controlled environmental conditions (55% ± 10% humidity, 22°C ± 2°C temperature, and a 12-h day/night cycle). Genotyping was performed at the IGBMC institute (Illkirch-Graffenstaden, France) using ear biopsies collected at weaning, following the genotyping protocols for *Apc^tm1Tno^, Kras^tm4Tyj^* and *Cre/ERT2* provided by the Jackson Laboratory. At eight to ten weeks of age, male and female mice were randomly allocated to experimental groups (n=16, 8 males and 8 females), based on genotype, and received two 10mg/kg intraperitoneal administrations of tamoxifen (Sigma-Aldrich, St. Louis, MO, USA) at 24 hours interval. Animal protocols and procedures were reviewed and approved by the French Nuclear Safety and Radiation Protection Authority Ethics Committee on Animal Experimentation and the French Ministry for Superior Education and Research (reference APAFIS #47235-2024020510357813 v2), and experiments were conducted in compliance with the European and French regulations on animal use for scientific purposes (EC Directive 2010/63/EU and French Decree 2013-118).

### Animal irradiations

Mouse irradiations were performed using a Small Animal Radiation Research Platform (SARRP, Xstrahl, Walsal, UK). Irradiations were conducted at 120 kV with a tube current of 5.0-5.2 mA and 0.8 mm beryllium and 0.15 mm copper filtration. A custom-designed collimator was used to generate a 4×8 cm irradiation field, enabling low-dose radiation exposure. Before each irradiation session, dosimetry verification was performed using a PTW3010 ionization chamber calibrated to water dose. The chamber was inserted into a mouse dosimetry phantom under conditions replicating those used for live-animal irradiation. Mice were placed in the supine position so that the beam entered through the abdominal region and were irradiated individually under gas anesthesia (2.5% isoflurane). Animals received single exposures of 25 mGy or 250 mGy (dose to water). To assess the impact of radiation at distinct stages of carcinogenesis, two irradiation schedules were implemented. Early irradiation was performed one week after tamoxifen administration, corresponding to the initial phase of tumor initiation. Late irradiation was delivered four weeks before euthanasia to assess radiation effects during tumor progression.

### Tissue collection

Animals were euthanized by cervical dislocation under anesthesia. For Apc^fl/fl^ Kras^fl/+^ and Apc^fl/fl^ Kras^+/+^ Cre^+^ mice (respectively referred to as [Control] and [Apc] mice), tissue collection was performed 12 weeks after tamoxifen administration. For Apc^fl/fl^ Kras^fl/+^ Cre^ERT2+^ mice (referred to as [ApcKras] mice), sampling was conducted 4-8 weeks after tamoxifen administration. Colons were harvested and processed using the improved Swiss-rolling technique [14]. Feces were collected directly from the colons and immediately frozen in liquid nitrogen.

### Histological preparation and analysis

Colons were fixed in 4% formaldehyde for a minimum of 24 hours, followed by standard dehydration and paraffin embedding. Five-micrometer sections were stained with hematoxylin, eosin and safranin (HES). The entire length of each colon was systematically annotated under the guidance of a trained pathologist, following criteria adapted from Boivin *et al.* [15]. The lesions were characterized as follows: normal (intact mucosal architecture, without histological abnormalities), hyperplasia (presence of at least two of the following features: crypt elongation exceeding 50% of normal length, increased number of mitotic figures, scalloped or irregular luminal contour), low-grade dysplasia (mild architectural distortion with loss of enterocyte alignment and occasional mucosal branching) or high-grade dysplasia (pronounced cellular atypia and marked architectural disorganization). Representatives images are provided in Fig. S1.

### L-WRN culture

L-WRN fibroblast-like cells (ATCC CRL-3276) were culture according to ATCC recommendations to produce Wnt-3A, R-spondin and Noggin conditioned medium. Cells were maintained in DMEM (Thermo Fisher Scientific, Waltham, MA, USA) supplemented with 10% FBS (Gibco, Carlsbad, CA, USA), and selective pressure was preserved using 0.5 mg/mL G- 418 (Gibco) and 0.5 mg/mL hygromycin B (Gibco). Conditioned medium was collected daily for three weeks following confluence. G-418 and hygromycin B were omitted during the collection period. The collected medium was pooled and stored at -20°C until use.

### Organoid generation

KPC:APC male mice were euthanized by carbon dioxide inhalation, after which colons were harvested and cut into small fragments. Tissue pieces were incubated at 37°C for 35 min in 10 mM EDTA (Thermo Fisher Scientific) prepared in PBS to release colonic crypts. Approximately 150 crypts were embedded in 30µL droplets of growth factor-reduced Matrigel (Corning, Corning, NY, USA) supplemented with 1x B27 (Thermo Fisher Scientific), 1x N2 (Thermo Fisher Scientific), 1µM N-acetylcysteine (Thermo Fisher Scientific), 50 ng/mL EGF (Thermo Fisher Scientific) and 10µM Y-27632 (BD Biosciences, Franklin Lakes, NJ, USA). A single 30µL Matrigel droplet was plated per well in a 24-well plate. The complete organoid culture medium consisted of advanced DMEM/F12 (Thermo Fisher Scientific) supplemented with 50% L-WRN conditioned medium, 1x N2 (Thermo Fisher Scientific), 1 µM N-acetylcysteine (Thermo Fisher Scientific), 50 ng/mL EGF (Thermo Fisher Scientific), 2 mM Glutamax (Thermo Fisher Scientific) and 5% FBS (Gibco). Medium was changed every 2-4 days. For the first two days of culture, 0.1mg/mL primocine (InvivoGen, San Diego, CA, USA) and 10µM Y-27632 (BD Biosciences) were added. Organoids were maintained at 37°C in a humidified incubator with 5% CO_2._

### Organoid treatment with 4-OHT

On day 3 of organoid culture, the original culture medium was replaced with complete culture medium supplemented with 0.125µM 4-hydroxytamoxifen (4-OHT, Merck, Darmstadt, Germany) and 10µM Y-27632 (BD Biosciences). Organoids were incubated under these conditions for 24 hours to induce Cre-mediated recombination.

### Organoid irradiation

Colon organoids were irradiated at doses of 25 and 250mGy, four days after initial seeding, using the same SARRP irradiation setup as for the *in vivo* animal experiments but without the mouse-specific collimator. A dosimetric verification was conducted prior to each irradiation session using a PTW31010 ionization chamber calibrated to air dose and positioned in a standard cell culture plate. Immediately after irradiation, organoids were returned to standard culture conditions.

### Organoid passage

Organoids were passaged seven days after culture initiation. Organoids were dissociated by incubating the cultures in TrypLE Express (Thermo Fisher Scientific) for 10 min at 37 °C. Resulting cell clusters and fragments were washed sequentially with PBS and washing medium composed of DMEM/F12 (Thermo Fisher Scientific), 10% FBS (Gibco), L-glutamine (Merck), and Penicillin–Streptomycin (Thermo Fisher Scientific). Cells were then resuspended in fresh supplemented Matrigel and replated as described above. The splitting ratio was adjusted based on organoid genotype and pre-passage density. Complete culture medium was added after polymerization of the Matrigel and refreshed every 2–4 days. For the first three days after passaging, 10 µM Y-27632 (BD Biosciences) was included to enhance post-dissociation survival. Organoids were collected for analyses (RNA and protein extraction) seven days after passaging. Live imaging and growth monitoring were performed using an IncuCyte S3 system (Sartorius). Analyses were performed on independent organoid cultures, derived from separate mouse individuals (n=2 for model validation, n=3-4 for radiation effects evaluation).

### EdU immunostaining

For proliferation analysis, organoids were cultured in 8-well Nunc Lab-Tek chamber slides (Thermo Fisher Scientific) following passaging. EdU incorporation was assessed using the Click-iT™ EdU Cell Proliferation Kit for Imaging (Thermo Fisher Scientific) according to the manufacturer’s instructions. Briefly, organoids were incubated with 10 µM EdU in complete culture medium for 1 hour at 37 °C, followed by fixation in 4% paraformaldehyde. After permeabilization, the Click-iT reaction cocktail was applied to visualize EdU-positive nuclei. Slides were mounted using 0.5 mm imaging spacers (Invitrogen) and a mounting medium containing DAPI to counterstain nuclei. Confocal imaging was performed using a Zeiss LSM 780 microscope. Image analysis was performed on Imaris 11 with the “Spots model” tool and the following parameters: objects estimated XY diameter = 6,00 µm, true background subtraction, filter spots quality and intensity center.

### RNA sequencing (RNA-seq)

RNA was extracted using the RNeasy Plus Kit (QIAGEN) following the manufacturer instructions. mRNA was selected by polyA-selection, then sequencing libraries were prepared using the NEBNext UltraTM RNA Library Prep Kit for Illumina (New England Biolabs, Ipswich, MA, USA) according to the manufacturer’s instructions. Paired-end RNA-seq was performed at BMK Gene (Münster, Germany) on a NovaSeq 6000 machine (Illumina, San Diego, CA, USA) to generate at least 20 million reads. Adaptor sequences and low-quality sequence reads were removed, then the clean reads were mapped to the Ensembl GRCm38 mouse genome using HISAT2 [16] and assembled using StringTie [17] using the recommended workflow (assembled transcripts were merged with the --merge option to generate a non-redundant set of transcripts, then a read coverage table was generated with the --eB option). Analysis of differential gene expression was performed with the DESeq2 R package using the median of ratios normalization method [18].

### m^6^A methylated RNA immunoprecipitation sequencing (meRIP-seq)

RNA was fragmented then divided into two aliquots: one for the m6A-inmunoprecipitation procedure (IP sample), and the other to be used as a non-inmunoprecipitated input control (input sample). The IP sample was incubated with an antibody-bead mixture, containing the anti-m6A antibody (Synaptic Systems, Göttingen, Germany), the protein-A and protein-G magnetic beads (Thermo Fisher Scientific, Waltham, MA, USA), and RNasin Plus RNase Inhibitor (Promega, Madison, WI, USA) for 2 h at 4°C. After extensive washing, the m6A-enriched RNA was purified using the RNeasy MiniElute spin column (Qiagen, Venlo, Netherlands). For library preparation, the SMARTer Stranded Total RNA-Seq Kit v2 - Pico Input Mammalian (Takara Bio, Kusatsu, Japan) was used following manufacturer’s instructions. Samples were dual-indexed for post-sequencing demultiplexing. Equimolar amounts of the libraries were sequenced at AllGenetics & Biology SL (A Coruña, Spain) using a NovaSeq PE150 flow cell (Illumina), aiming for a total output of 155 Gb. The raw sequencing data were mapped to the GRCm39 mouse reference genome by STAR aligner [19] and the resulting BAM files were analyzed using the “RNA methylation Differential Analysis in R” (RADAR) R package [20].

### Cell lysis and protein quantification

Proteins were extracted using RIPA lysis buffer (Thermo Fisher Scientific) supplemented with protease inhibitors (cOmplete Tablets, Roche, Basel, Switzerland). Lysates were incubated on ice and clarified by centrifugation. Total protein concentration was determined using the Pierce Rapid Gold BCA Protein Assay Kit (Thermo Fisher Scientific), following the manufacturer’s instructions.

### Mass spectrometry proteomics

Preparation and analysis of samples for proteomic experiments were performed at the Proteogen platform of Caen University (France). Protein extracts were prepared using a modified Gel-aided Sample Preparation protocol [21]. Samples were digested with trypsin/Lys-C overnight at 37°C. For nano-LC fragmentation, peptide samples were first desalted and concentrated onto a µC18 Omix (Agilent) before analysis. The chromatography step was performed on a NanoElute (Bruker Daltonics) ultra-high-pressure nano flow chromatography system. Approximatively 50ng of each peptide sample were concentrated onto a C18 pepmap 100 (5mm x 300µm i.d.) precolumn (Thermo Scientific) and separated at 50°C onto a reversed phase Reprosil column (25cm x 75μm i.d.) packed with 1.6μm C18 coated porous silica beads (Ionopticks). Mobile phases consisted of 0.1% formic acid, 99.9% water (v/v) (A) and 0.1% formic acid in 99.9% ACN (v/v) (B). The nanoflow rate was set at 250 nl/min, and the gradient profile was as follows: from 2 to 15% B within 15 min, followed by an increase to 30% B within 15 min, to 45% B in 5min and further to 95% within 5 min and reequilibration. MS experiments were carried out on an TIMS-TOF HT mass spectrometer (Bruker Daltonics) with a modified nano electrospray ion source (CaptiveSpray, Bruker Daltonics). A 1400 spray voltage with a capillary temperature of 180°C was typically employed for ionizing. MS spectra were acquired in the positive mode in the mass range from 100 to 1700 m/z and 0.75 to 1.28 1/k0 window. In the experiments described here, the mass spectrometer was operated in PASEF DIA mode with exclusion of single charged peptides. The DIA acquisition scheme consisted of 24 variable windows ranging from 350 to 1000 m/z.

Database searching and label free quantification (using XIC) was performed using DIA-NN (version 2.0) [22]. An updated *Mus musculus* database was used for library-free search / library generation. For retention time prediction and extraction mass accuracy, we used the default parameter 0.0, which means DIA-NN performed automatic mass and retention time correction. Top six fragments (ranked by their library intensities) were used for peptide identification and quantification. The variable modifications allowed were as follows: Nterm-acetylation and Oxidation (M). “Trypsin/P” was selected. Data were filtering according to a FDR of 1%. Cross-run normalisation was performed. To quantify the relative levels of protein abundance between different groups, data from DIA-NN were then analysed using a modified DEP package from R. Briefly, proteins that are identified in 2 out of 3 replicates of at least one condition were filtered, missing data were imputed using a random draw from a Gaussian distribution centered around a minimal value and differential enrichment analysis was based on linear models and empirical Bayes statistic. A 1.2-fold increase in relative abundance and a 0.05 p-value were used to determine enriched proteins.

### Enrichment analysis

Cancer-related genes and proteins were selected and annotated as oncogenes or tumor suppressors based on the COSMIC Cancer Gene Census [23] and the OncoKB cancer gene database [24]. Their contribution to carcinogenesis was determined by integrating their functional classification (oncogene vs. tumor suppressor) with their observed direction of dysregulation in the datasets. Enrichment in cancer hallmarks was evaluated using the “CancerHallmarks” online analysis tool [25]. Gene set enrichment analysis (GSEA) was performed using the GSEA 4.3.2 software with gene sets derived from the MSigDB v2025.1.Mm collection [26–28].

### Western Blot

A total of 30µg of protein lysate was denatured at 37°C for 30 min in sample buffer containing NuPAGE LDS Sample buffer (Thermos Fisher Scientific) and NuPAGE Sample Reducing Agent ((Thermos Fisher Scientific). Samples were resolved on 4-10% polyacrylamide gels (Mini-protean TGX, BioRad) with MES buffer (MES SDS NuPAGE, Invitrogen) and transferred on PVDF membranes (Trans-Blot Turbo Transfert Pack PVDF Mini, BioRad). Aspecific sites were blocked with 1 hour, 5% BSA incubation before overnight incubation with primary antibodies: anti-mTOR antibody (Abcam ab25880), anti-mTOR phospho S2448 antibody (abcam, ab131538), anti-phospho-4E-BP1 Thr37/46 (Cell signaling, 2855). The proteins were visualized after anti-rabbit-HRP (R&D systems, HAF008) incubation by chemiluminescence with the Imagequant 800 (Cytiva). The membranes were stripped with Restore^TM^ Western Blot stripping buffer (Thermo Scientific) and incubated with a second batch of primary antibodies: anti-4E-BP1 (Cell signaling, 9644). GAPDH was used for protein normalization.

### Statistical analysis

Statistical analysis was performed on GraphPad Prism 10.1.1.323 or with the GSEA 4.3.2 software. Statistical analyses on the distribution of pre-cancerous lesions were performed using the chi² test. Multiple comparisons of irradiated conditions against NIR were performed with Dunnett’s test, while all pairwise comparisons were performed using Tukey’s test. Non-parametric statistical evaluation was conducted with the Kruskal-Wallis test. Statistical significance was considered when p-value was < 0.05. For omics analyses, significance was assessed via False Discovery Rate for differentially expressed (DE) genes and GSEA. Significance was considered when FDR < 0.05 for DE genes and proteins, 0.1 for GSEA analysis.

## 3. Results

### Radiation exposure modulates carcinogenesis in Kras-mutated mice

The impact of medical diagnostic-range radiation (25 mGy, representative of a single abdominal CT examination, and 250 mGy, representative of cumulative CT exposures) on colon carcinogenesis was assessed by quantifying epithelial areas exhibiting hyperplasia, low-grade dysplasia, and high-grade dysplasia. Unsurprisingly, tamoxifen administration did not affect the colonic epithelium in unmutated mice (*Apc*^fl/fl^ *Kras*^fl/+^, hereafter [Control]) (Fig. 1A), whereas it led to a decrease in the normal epithelium fraction in *Apc*-mutated mice (*Apc*^fl/fl^ *Kras*^+/+^ Cre^ERT2+^, hereafter [Apc]) and in mice harboring both *Apc* and *Kras* mutations (Apc^fl/fl^ Kras^fl/+^ Cre^ERT2+^, hereafter [ApcKras]) (Fig. 1B-D). As no sex-specific differences were observed in lesion distribution in [Control] or [Apc] mice (Fig. S2A-D), male and female mice were pooled for analysis in these genotypes. For both groups, exposure to 25 or 250 mGy did not modify the development of pre-cancerous lesions, regardless of irradiation timing (Fig. 1A-B). In contrast, [ApcKras] mice displayed sex-specific responses to radiation. Although no difference in lesion distribution was detected between males and females under non-irradiated (NIR) conditions, significant sex-dependent effects emerged following exposure to LDR or moderate dose radiation (MDR) (Fig. S2E-F). Consequently, data from male and female [ApcKras] mice were analyzed separately.

**Figure 1:**
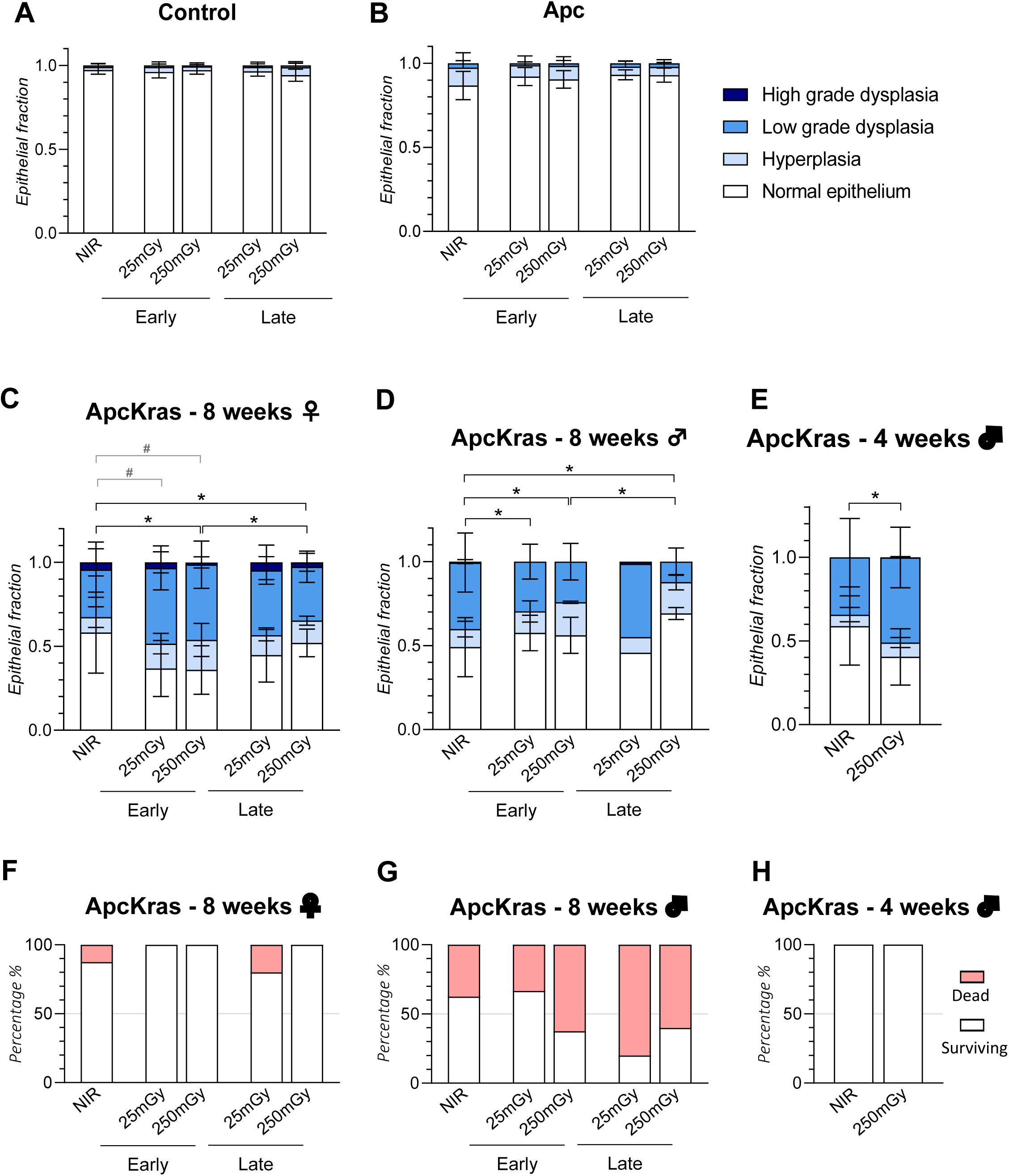
Pre-cancerous lesions can be observed in the colonic epithelium of KPC:APC mice after tamoxifen administration and irradiation. Epithelial fractions presenting pre-cancerous lesions (as a percentage of the whole colonic epithelium area) for: A) [Control] mice (male and female) 12 weeks after tamoxifen treatment; B) [Apc] mice (male and female) 12 weeks after tamoxifen treatment; C) [ApcKras] female mice 8 weeks after tamoxifen treatment, significant chi² test performed on lesion distribution is represented using the “*” symbol, significant Tukey’s test performed on total abnormal areas is represented using the “#” symbol, D) [ApcKras] male mice 8 weeks after tamoxifen treatment, E) [ApcKras] male mice 4 weeks after tamoxifen treatment and exposed to early radiation, F) Percentage of surviving mice on [ApcKras] female mice 8 weeks after tamoxifen treatment, n=5-8, G) Percentage of surviving mice on [ApcKras] male mice 8 weeks after tamoxifen treatment, n=5-8 and H) Percentage of surviving mice on [ApcKras] male mice 4 weeks after tamoxifen treatment and irradiated early, n=4.

In female [ApcKras] mice, lesion distribution was significantly altered following both early and late irradiation at 250 mGy, with a more pronounced effect in the early-irradiation group (Fig. 1C). After early irradiation (performed 1 week after tamoxifen-induced carcinogenesis initiation), the fraction of normal epithelium decreased significantly, with no evidence of dose dependence. Concurrently, areas of low-grade dysplasia increased significantly following exposure to 25 and 250 mGy. On the contrary, after late irradiation (4 weeks prior to euthanasia), the changes in normal and abnormal epithelium fraction area were not significant. Male [ApcKras] mice exhibited an opposite trend (Fig. 1D). Following irradiation, the combined area of hyperplasia, low-grade dysplasia, and high-grade dysplasia was slightly reduced compared to controls.

However, these findings must be interpreted with caution, as mortality exceeded 50% in several irradiated male groups, potentially introducing bias in lesion distribution (Fig.1F-H). To evaluate this possibility, we analyzed a separate cohort of [ApcKras] male mice collected 4 weeks after tamoxifen administration and exposed to early irradiation (250 mGy). This analysis confirmed a significant reduction in the fraction of normal epithelium following irradiation (Fig. 1E). A small number of [ApcKras] animals developed colonic adenomas. Although the number of adenomas per mouse was too low to assess changes in multiplicity following irradiation, adenoma size was quantified (Fig. S2G). Female mice exhibited a non-significant increase in adenoma size after early 250 mGy exposure. While male and female [ApcKras] mice developed a comparable number of adenomas under NIR conditions (Figure S2H), no adenomas were detected in irradiated male groups. Notably, under NIR conditions, adenomas in male mice were significantly larger than those observed in females.

Because the gut microbiota plays a critical role in intestinal physiology and CC development, we also performed metagenomic sequencing of fecal samples from experimental mice [29]. Analysis of the gut microbiota in irradiated and non-irradiated [Control] and [ApcKras] mice revealed no significant changes in the abundance of pathogenic or cancer-associated bacterial genera. Irradiation induced a decrease in alpha diversity exclusively in [Control] mice, whereas no diversity changes were detected in [ApcKras] animals (Fig. S3).

### Radiation exposure modulates organoid transformation

To investigate the mechanisms underlying the response to diagnostic radiation, KPC:APC organoids were generated from KPC:APC mice. This dynamic *ex vivo* model recapitulates key steps of colon carcinogenesis through the *in vitro* induction of *Apc* and *Kras* mutations. Organoids derived from [Apc] and [ApcKras] mice displayed phenotypic and molecular features characteristic of the carcinogenic-like process, including a pronounced cystic morphology (Fig. 2A-B), increased organoid size (Fig. 2A and 2C), enrichment of cancer-associated pathways at both transcriptomic and proteomic levels compared with [Control] organoids (Fig. 2D-E), and distinct proteomic signatures (Fig. 2F). The organoid model also faithfully captures changes in m6A RNA methylation associated with colorectal cancer progression (Fig. 2G and 2G, Fig. S4). Among all genotypes and irradiation conditions, [ApcKras] organoids exposed early to 250 mGy were the only group exhibiting a significant increase in size following irradiation, with their average measured area doubling compared with non-irradiated counterparts (Fig. 3A-C). Despite this increase in size, their overall morphology remained unchanged: the proportion of cyst-like versus differentiated organoids was not modified by early exposure (Fig. 3D). Proliferation analysis in [ApcKras] organoids revealed a significant increase specifically within the spheroid subpopulation (Fig. 3E-G).

**Figure 2:**
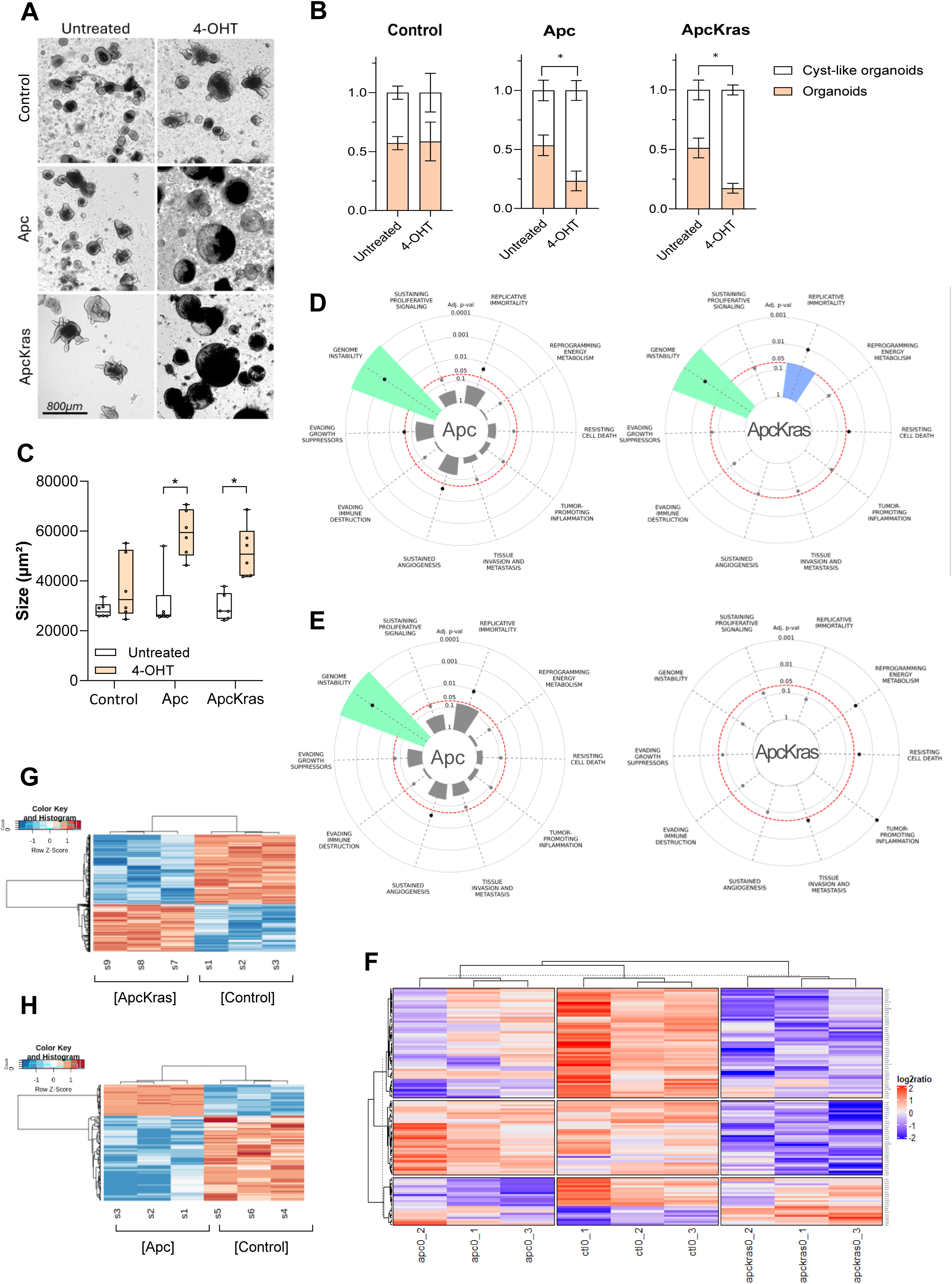
4-OHT administration triggers morphological and molecular changes in KPC:APC organoids. A) Representative images of organoids after 4-OHT administration. B) Organoid morphology ratios (cyst-like vs budding-type differentiated organoids). C) Organoid size after 4-OHT administration. Each point represents the mean organoid size per well from 3 independent experiments. D) Overrepresentation of hallmark-associated genes among DE genes. E) Overrepresentation of hallmark-associated proteins among differentially expressed (DE) proteins. F) Hierarchical clustering of DE proteins across genotypes. G-H) Heatmap of differentially methylated (DM) transcripts in [Control] and [ApcKras] (G) or [Apc] (H) organoids.

**Figure 3:**
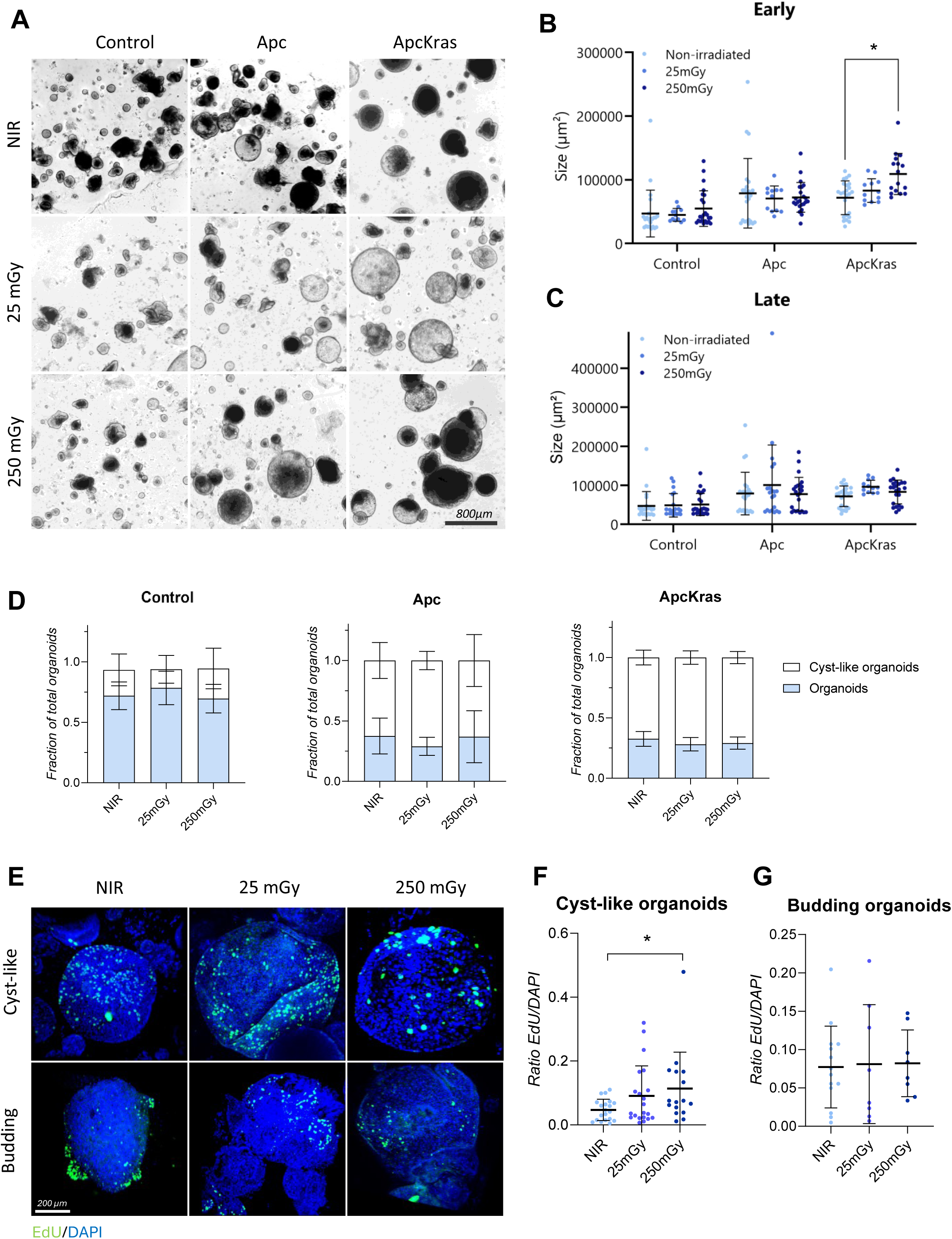
Diagnostic radiation induces morphological changes and stimulates the proliferation of KPC:APC organoids. A) Representative images of KPC:APC organoids following diagnostic radiation exposure. B) Organoid average size variation following early exposure. C) Organoid average size variation following late exposure. D) Fractions of cyst-like and normal organoids in [Control], [Apc] and [ApcKras] organoids. E) Representative images of EdU labeling with classification of cyst-like and budding organoids. F-G) Proliferation assessment of [ApcKras] cyst-like (F) or budding (H) organoids by quantification of EdU labeling.

### Medical diagnostic radiation triggers cancer-associated transcriptome alterations

The response to diagnostic radiation was first evaluated at the transcriptomic level by RNA sequencing. The number of differentially expressed (DE) genes varied according to genotype and radiation dose (Fig. 4A). Exposure to 250 mGy induced a broader transcriptional dysregulation than 25 mGy in both [Apc] and [ApcKras] organoids, with the [ApcKras]–250 mGy condition presenting the highest numbers of both up- and down-regulated genes. Notably, only a small subset of DE genes overlapped between the responses to 25 mGy and 250 mGy, indicating largely dose-specific transcriptional signatures (Figure 4B).

**Figure 4:**
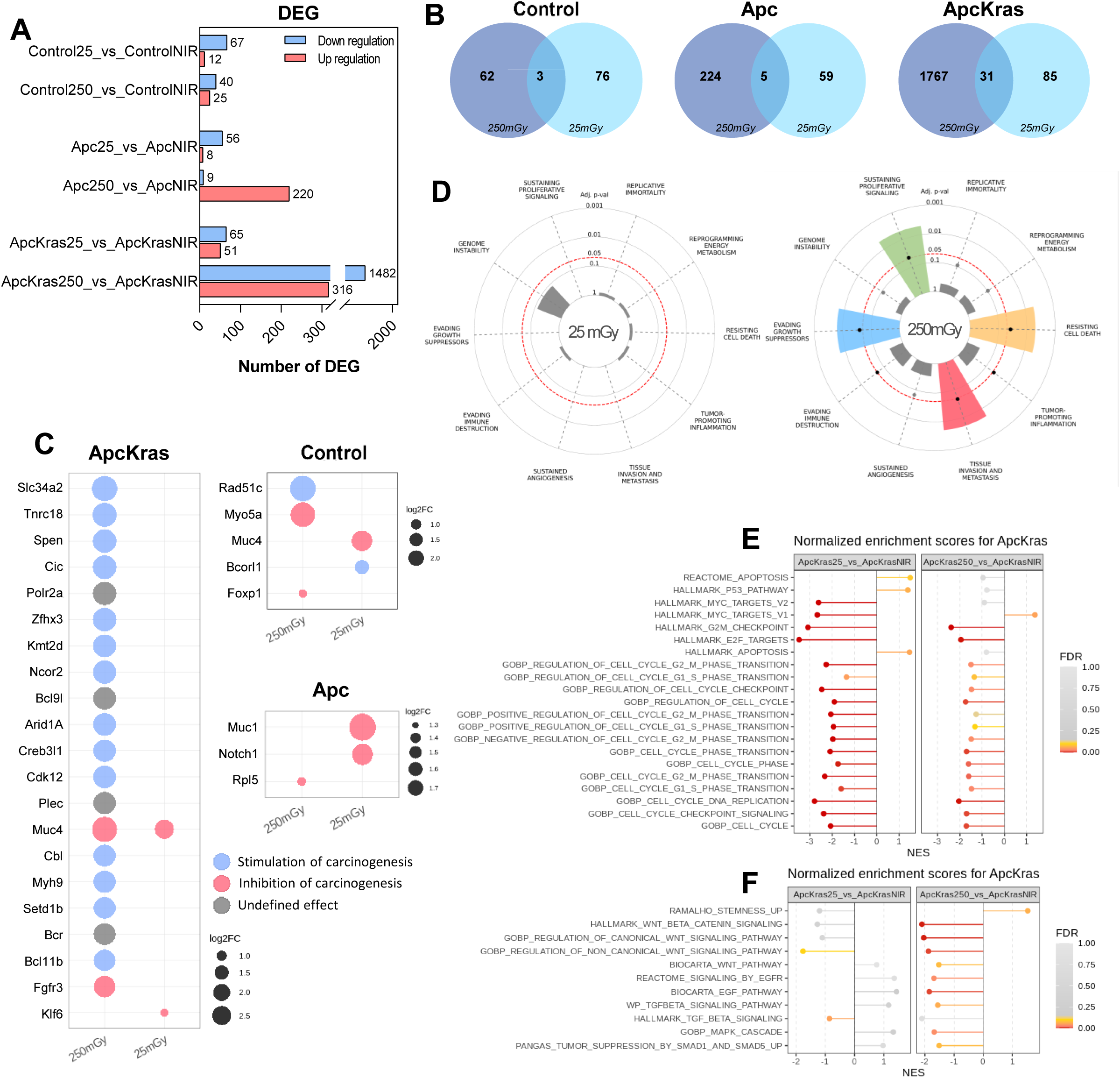
Diagnostic radiation triggers cancer-associated transcriptomic changes in KPC:APC organoids. A) DE genes (FC>1.5, adjusted p-value <0.05). B) Venn diagram showing overlap of DE genes per genotype. C) Impact of the top 20 differentially expressed cancer-related genes dysregulation on carcinogenesis. Tumor suppressor genes are indicated in pink and oncogenes in blue. Genes whose role as oncogene or tumor suppressor is unclear or subject to variation are indicated in black. D) Overrepresented cancer hallmarks in irradiated [ApcKras] compared to non-irradiated. E) GSEA enrichment in proliferation and cell cycle pathways in [ApcKras]. Gene sets included in the figure had an FDR<0.1 in at least one comparison. F) GSEA enrichment in colon cancer-related pathways in [ApcKras] organoids. Gene sets included in the figure had an FDR<0.1 in at least one comparison.

To determine whether radiation-induced transcriptional changes were functionally linked to carcinogenesis, the 20 most differentially expressed cancer-related genes in each condition were examined (Fig. 4C). [ApcKras] organoids irradiated at 250 mGy were the only group exhibiting at least 20 dysregulated cancer-associated genes compared with their NIR counterparts. These genes predominantly promote carcinogenesis-like features through down-regulation of tumor suppressor genes. To further explore which cancer-associated functions were altered by irradiation, we assessed the overrepresentation of hallmark-specific genes within the DE gene lists from [ApcKras] organoids. A significant enrichment of multiple cancer hallmarks was observed after 250 mGy exposure, including pathways related to sustaining proliferative signaling, evading growth suppression, resisting cell death, and tissue invasion and metastasis (Figure 4D).

To elucidate the pathways and molecular mechanisms perturbed by diagnostic radiation, gene set enrichment analysis (GSEA) was performed using gene sets associated with cell proliferation (Fig. 4E) and CC (Fig. 4F). [ApcKras] organoids demonstrated strong enrichment for proliferation-related gene sets, including under LDR conditions (25 mGy). Overall, numerous cell-cycle-associated gene sets exhibited negative enrichment scores, consistent with perturbations in cell-cycle regulation. Enrichment of CC–related pathways was primarily observed after MDR exposure. Except for stemness-associated gene sets, most CC–related pathways displayed negative enrichment, with the strongest changes involving Wnt-pathway regulation and EGF-signaling gene sets.

### Medical diagnostic radiation triggers cancer-relevant proteomic changes

The proteomic response to diagnostic radiation in organoids was assessed by mass spectrometry. As observed at the transcriptomic level, exposure to 250 mGy induced a greater number of DE proteins than 25 mGy across all genotypes (Fig. 4A). However, in contrast to transcriptomic analyses, [ApcKras] organoids exhibited the fewest DE proteins overall. A larger fraction of DE proteins was shared between the two irradiation doses compared with the overlap observed among DE genes (Fig. 4B), indicating a more conserved dose-dependent signature at the protein level. Analysis of the 20 most dysregulated proteins encoded by oncogenes and tumor suppressors did not reveal a consistent pattern (Fig. S5). Although 250 mGy exposure induced larger fold-changes than 25 mGy across all genotypes, no clear trend toward predominantly pro- or anti-carcinogenic-like protein dysregulation emerged. Given the absence of clear directional patterns, enrichment of cancer-hallmark-associated proteins was also examined. Only [ApcKras] organoids demonstrated a significant overrepresentation of proteins linked to genome instability following 250 mGy exposure (Fig. 4C; Fig. S6A and S6B), suggesting a genotype-specific proteomic response to diagnostic radiation.

To directly compare proteomic and transcriptomic effects, gene-set enrichment analyses focused on proliferation and cancer-related pathways were performed. Similar to transcriptomic findings, the strongest and most significant enrichment was observed following 25 mGy exposure (Fig. 4D). Many enriched sets corresponded to regulators of the cell cycle, particularly those involved in phase-transition control. CC–related pathways showed consistent enrichment patterns, with Mammalian target of rapamycin complex 1 (mTORC1) signaling emerging as the only pathway significantly and positively enriched at both 25 and 250 mGy (Fig. 5E and 5F). This finding highlights mTORC1 as a shared potential effector of diagnostic radiation across doses. Western blot quantification of mTOR, Ribosomal protein S6 kinase (S6K) and Eukaryotic translation initiation factor 4E-binding protein 1 (4E-BP) activation suggested possible activation of S6K and 4E-BP signaling, the main targets of mTORC1, even though increases were not significant due to interexperimental variability and small number of replicates (Fig. 4G-H, Fig. S7).

**Figure 5:**
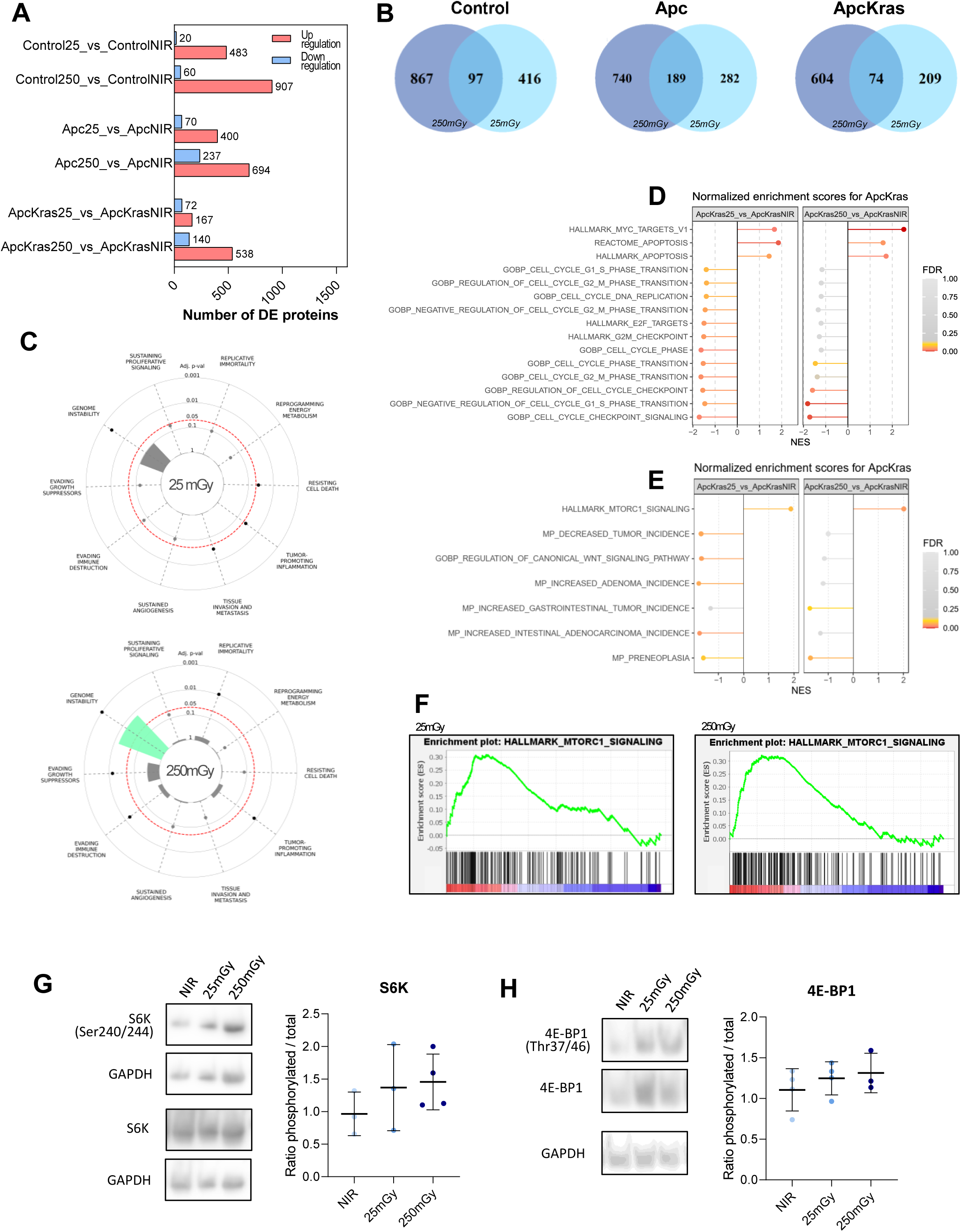
Diagnostic radiation induces cancer-associated proteomic changes in KPC:APC organoids. A) Differentially expressed (DE) proteins (FC>1.2, adjusted p-value<0.05). B) Venn diagram showing overlapping DE proteins per genotype C) Overrepresented cancer hallmarks in irradiated [ApcKras] compared to non-irradiated. D) GSEA enrichment in proliferation and cell cycle pathways in [ApcKras]. E) GSEA enrichment in colon cancer-related pathways in [ApcKras]. Gene sets included in the figure had an FDR<0.1 in at least one comparison. F) mTORC1 signaling enrichment plots in [ApcKras] organoids. G) Evaluation of S6K phosphorylation in [ApcKras] organoids. H) Evaluation of 4E-BP phosphorylation in [ApcKras] organoids.

### Medical diagnostic radiation induces cancer-associated m^6^A RNA methylation changes

Because m^6^A RNA methylation has recently been to be implicated in cellular responses to various genotoxic and non-genotoxic stresses, including exposure to IR [30], we assessed m^6^A methylation changes in irradiated KPC:APC organoids using methylated RNA immunoprecipitation sequencing (meRIP-seq). Following exposure to 250 mGy, the number of differentially methylated (DM) transcripts varied across genotypes (Table 1; Fig. 6A–5C). In [ApcKras] organoids, although the total number of DM transcripts was lower than in [Control] and [Apc] organoids, the top 1,000 DM bins clearly separated irradiated and non-irradiated samples by principal component analysis (Fig. 6D). Notably, approximately one-third of DM transcripts in [ApcKras] organoids (14/41) were associated with at least one canonical cancer hallmark. The hallmarks with the greatest number of DM transcripts included sustaining proliferative signaling, evading growth suppressors, tissue invasion and metastasis, and resisting cell death (Table S1). DM peaks were distributed across multiple transcript regions, including coding sequences (CDS), 3′ untranslated regions (3′ UTR), 5′ UTRs, and non-coding regions (Table S2). Altogether, these results suggest that the control of m6A RNA methylation might contribute to radiation-induced effects in [ApcKras] organoids.

**Figure 6:**
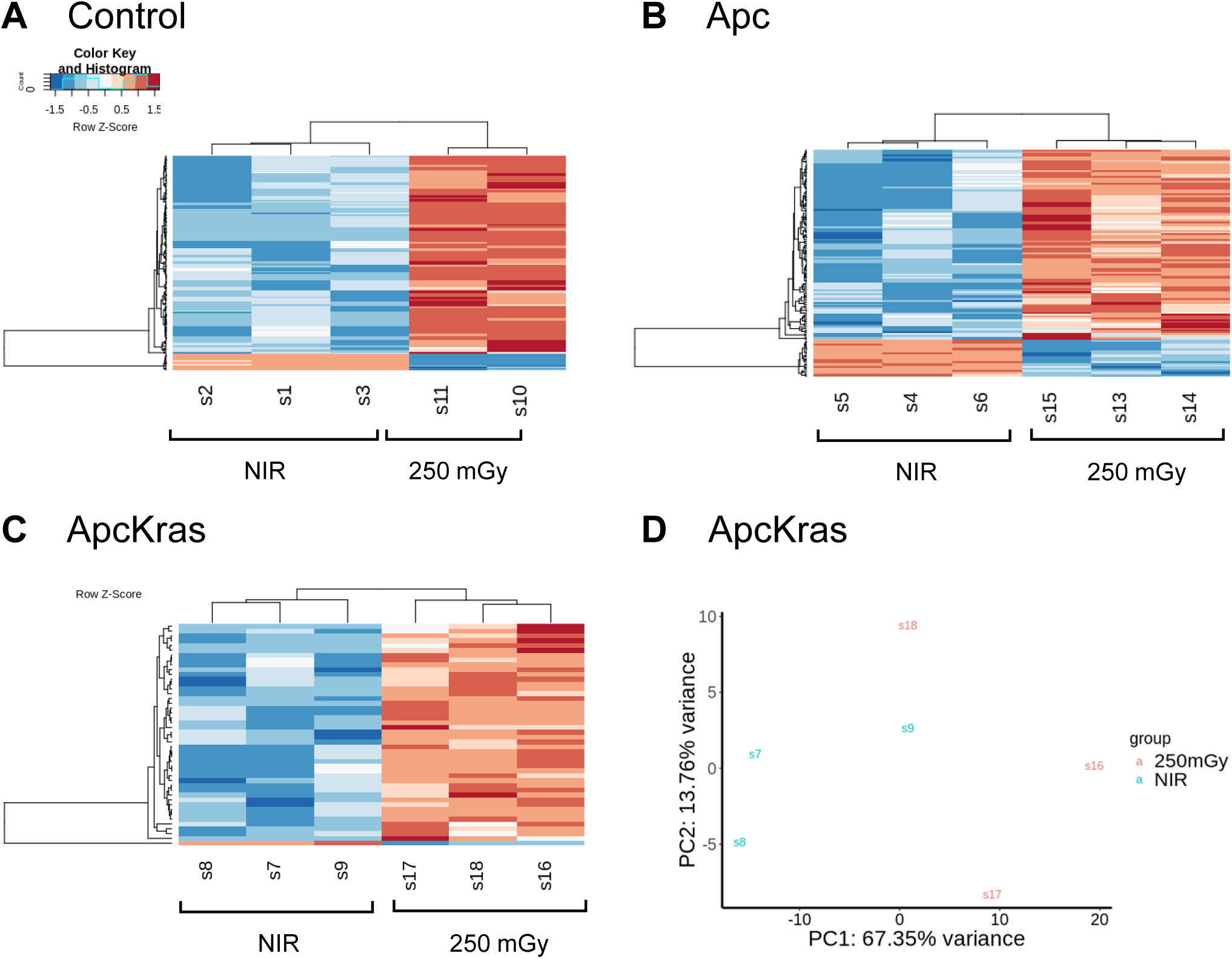
Diagnostic radiation modulates m^6^A RNA methylation in KPC:APC organoids. A-C) Heatmap of differentially methylated (DM) transcripts in [Control] (A), [Apc] (B) or [ApcKras] (C) organoids. D) Principal component analysis (PCA) on m6A methylation levels in the top 1,000 bins ranked by count number in [ApcKras] organoids.

**Table 1:**
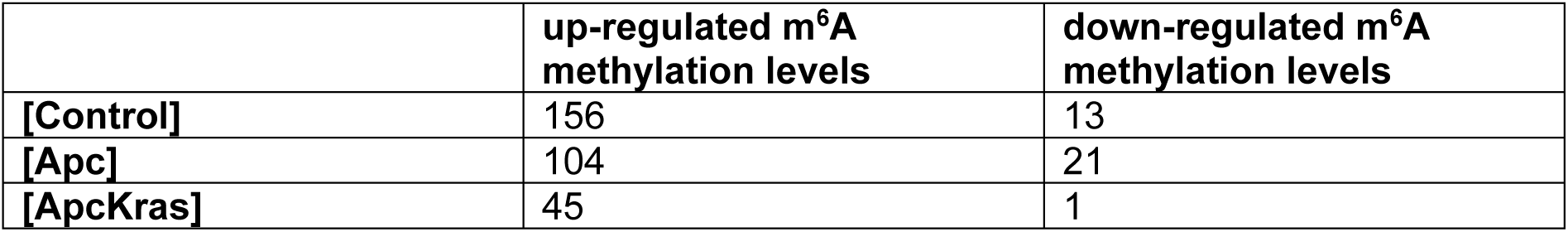
Number of differentially methylated (DM) transcripts after 250 mGy irradiation.

|  | up-regulated m<sup>6</sup>A methylation levels | down-regulated m<sup>6</sup>A methylation levels |
| --- | --- | --- |
| [Control] | 156 | 13 |
| [Apc] | 104 | 21 |
| [ApcKras] | 45 | 1 |

## 4. Discussion

Few studies have investigated the impact of medical diagnostic radiation on colon carcinogenesis, and existing findings remain inconclusive. Importantly, none have specifically addressed potential adverse effects associated with low-dose medical radiation. Using the KPC:APC mouse model, we demonstrate that exposure to 25 mGy (approximating a single CT scan) or 250 mGy (representing repeated CT scans) promotes the development of pre-cancerous lesions under defined genetic conditions. In mice lacking pre-disposing mutations, no radiation-induced carcinogenesis was detected. Likewise, in mice carrying the *Apc* mutation alone, diagnostic radiation did not accelerate lesion formation at either the tissue or molecular level—consistent with previous reports showing that *Apc* loss sensitizes colonic epithelium only to higher-dose irradiation (≥1 Gy) and with long latency (>6 months) [10,31]; similarly, chronic ɤ low-dose exposure did not affect tumor length and multiplicity in *Apc^Min/+^* mice [32]. In contrast, the combination of *Apc* and *Kras* mutations in our model rendered colonic epithelium markedly sensitive to diagnostic radiation.

Incidence and mortality from colorectal cancer are generally higher in men than in women [33]; investigations suggest that these differences might result from variations in sex hormone levels and the gut microbiota [34]. Similar differences were observed in mouse models of CC [35,36]. Surprisingly, radiation exposure appeared to exert opposite effects in male and female [ApcKras] mice, with males showing fewer or smaller lesions than females. However, this paradox is likely explained by survivor bias. Male [ApcKras] mice experienced substantial mortality 8 weeks after irradiation. Surviving males—those included in lesion quantification—had significantly larger adenomas under non-irradiated conditions (Fig. S2H), suggesting that more aggressive tumors may have contributed to early lethality. Consistent with this interpretation, no adenomas were detected in irradiated male groups; pre-cancerous lesion size increased 4 weeks post-exposure; and organoids derived from males exhibited radiation-stimulated carcinogenesis-like features.

Taken together, when accounting for survival-related selection, our data support the conclusion that diagnostic radiation enhances colon carcinogenesis in both male and female mice carrying *Apc* and *Kras* mutations, with no clear dose-dependent effect at the tissue level. The dependence of the response on genetic background is further supported by our omics analyses: the overlap in DE genes and proteins across irradiated genotypes was minimal (Fig. S8A and S8B), and only [ApcKras] organoids exhibited molecular signatures indicative of carcinogenesis-like stimulation. Moreover, both *in vivo* and organoid results suggest that radiation exposure during the initiation phase has a stronger impact than exposure during later progression.

To identify the molecular mechanisms driving radiation-enhanced carcinogenesis-like evolution, we examined gene and protein expression profiles in the novel KPC:APC organoids. [Apc] and [ApcKras] organoids presented a cyst-like phenotype characteristic of *Apc*-mutated organoid models [37,38] and molecular signatures validating their unique and carcinogenic-like status at p(1) +7 [39,40]. Both transcriptomic and proteomic analyses confirmed that oncogenic signaling was stimulated exclusively in [ApcKras] organoids. Proteomic profiling suggested mTORC1 signaling as a central pathway differentiating irradiated from non-irradiated samples, suggesting that mTORC1 activation may contribute to the pro-tumorigenic effects of radiation. This is consistent with the well-established role of mTORC1 in regulating cell proliferation, migration, invasion, epithelial–mesenchymal transition, and ultimately metastasis formation [41,42].

Indeed, cell proliferation was elevated in irradiated [ApcKras] organoids. mTORC1 promotes proliferation through translational control mechanisms, notably by phosphorylating 4E-BP1 [42]. Interestingly, the stimulation of proliferation was restricted to the cyst-like organoid subpopulation. Given that cyst-like morphology is associated with undifferentiated, tumor-like organoids [37], whereas budding structures reflect epithelial differentiation and polarization [43], these findings suggest that oncogenic transformation remains incomplete in budding [ApcKras] organoids. Selective promotion of proliferation in cyst-like organoids aligns with our primary observation that only advanced-stage double mutants exhibit diagnostic radiation sensitivity.

Part of this proliferative response may be mediated through c-Myc targets, which show dose-dependent enrichment at the proteomic level following exposure. c-Myc promotes proliferation via transcriptional activation and repression of genes regulating cell cycle progression and the G1/S transition [44,45]. Interestingly, mTORC1 activation enhances Myc translation by promoting degradation of Myc antagonists via S6K-dependent pathways [46,47]. This mechanistic provides a plausible explanation for the discordance observed in [ApcKras] organoids, in which Myc-related gene sets were negatively enriched at the transcriptomic level but positively enriched at the proteomic level. Although these data are consistent with activation of the mTORC1-S6K-Myc axis, further investigation is required to formally establish its involvement. Moreover, not all observed cell-cycle dysregulations can be attributed solely to mTORC1 signaling, indicating that additional molecular mechanisms likely contribute to the radiation response.

Investigating the molecular response at the transcriptomic and proteomic levels highlighted the contribution of post-transcriptional regulation to the radiation-associated dysregulation of colon cancer–related pathways. RNA methylation, particularly m6A modification, has recently been shown to mediate part of the cellular response to genotoxic and non-genotoxic stressors, including ionizing radiation [30,48], and is increasingly considered a potential biomarker of radiation [49]. In our study, exposure to 250 mGy did not induce global changes in m6A levels but instead resulted in the modulation of methylation of specific transcripts. It has recently been shown that both the position of m6A residues within the transcript (whether located in the coding sequence or untranslated regions) and the mRNA secondary structure determine whether m6A exerts a stimulatory or inhibitory effect on translation or mRNA decay [50,51]. In [ApcKras] organoids, several of the differentially methylated transcripts encode genes implicated in carcinogenesis, suggesting that m6A-dependent regulation may contribute to the observed effects of diagnostic radiation.

Notably, the cancer hallmarks most affected by differential m6A methylation were those whose enrichment was diminished when comparing transcriptomic and proteomic datasets, consistent with hallmark-specific post-transcriptional buffering. mTORC1 signaling may play an important role in this process, as it can stimulate m6A deposition and thereby reduce the translation of specific methylated transcripts [52]. However, these alterations alone do not explain the totality of the modifications observed nor the origin of the enrichment of the mTORC1 signaling pathway, suggesting the involvement of additional post-transcriptional mechanisms. Interestingly, cell cycle regulation exhibited strong concordance between transcriptomic and proteomic modulations (suggesting limited post-transcriptional regulation), except for Myc, whose alteration is consistent with its regulation by mTORC1 signaling as previously described [47].

While our organoid model is able to recapitulate key aspects of colon carcinogenesis *in vitro*, several limitations need to be considered. For instance, it cannot integrate effects due to the tumor microenvironment or the modulation of the gut microbiome, which are well-established modulators of CC development [53,54]. In our study, genotype-dependent alterations in microbial diversity were observed in irradiated KPC:APC mice, suggesting that microbiota changes may contribute to some of the radiation-associated effects, as proposed elsewhere [32]. Further investigations will be required to elucidate the contribution of non-mutational processes (including the interactions between tumor-initiating epithelial cells and tissue-resident immune populations) to colorectal cancer progression following irradiation.

Although the overall carcinogenic risk associated with medical diagnostic CT examinations is considered very low, our findings demonstrate that they may nonetheless promote carcinogenic development in predisposed cells. While CT imaging remains an essential diagnostic tool, these observations underscore the importance of integrating individual susceptibility into benefit-risk evaluations and of continuing efforts to minimize unnecessary radiation exposure through adherence to ALARA (“as low as reasonably achievable”) principles and optimization of imaging protocols.

## Data availability statement

Datasets are available in the ASNR institutional repository at recherche.data.gouv.fr https://doi.org/10.57745/ZYPM9S.

## Acknowledgements

The authors thank the ASNR animal care team, including Delphine Denais-Lalieve, Romain Granger and Frédéric Voyer. They also thank Morgane dos Santos for her help with SARRP setup and irradiation experiments, Véronique Ménart for SARRP organoid irradiations, and the Proteogen platform of the University of Caen Normandy, including Benoit Bernay, for their help with proteomics experiments.

