## Supplementary Information for "Medical diagnostic radiation promotes murine Apc/Kras-driven colon carcinogenesis"

#### 1. Supplementary figures

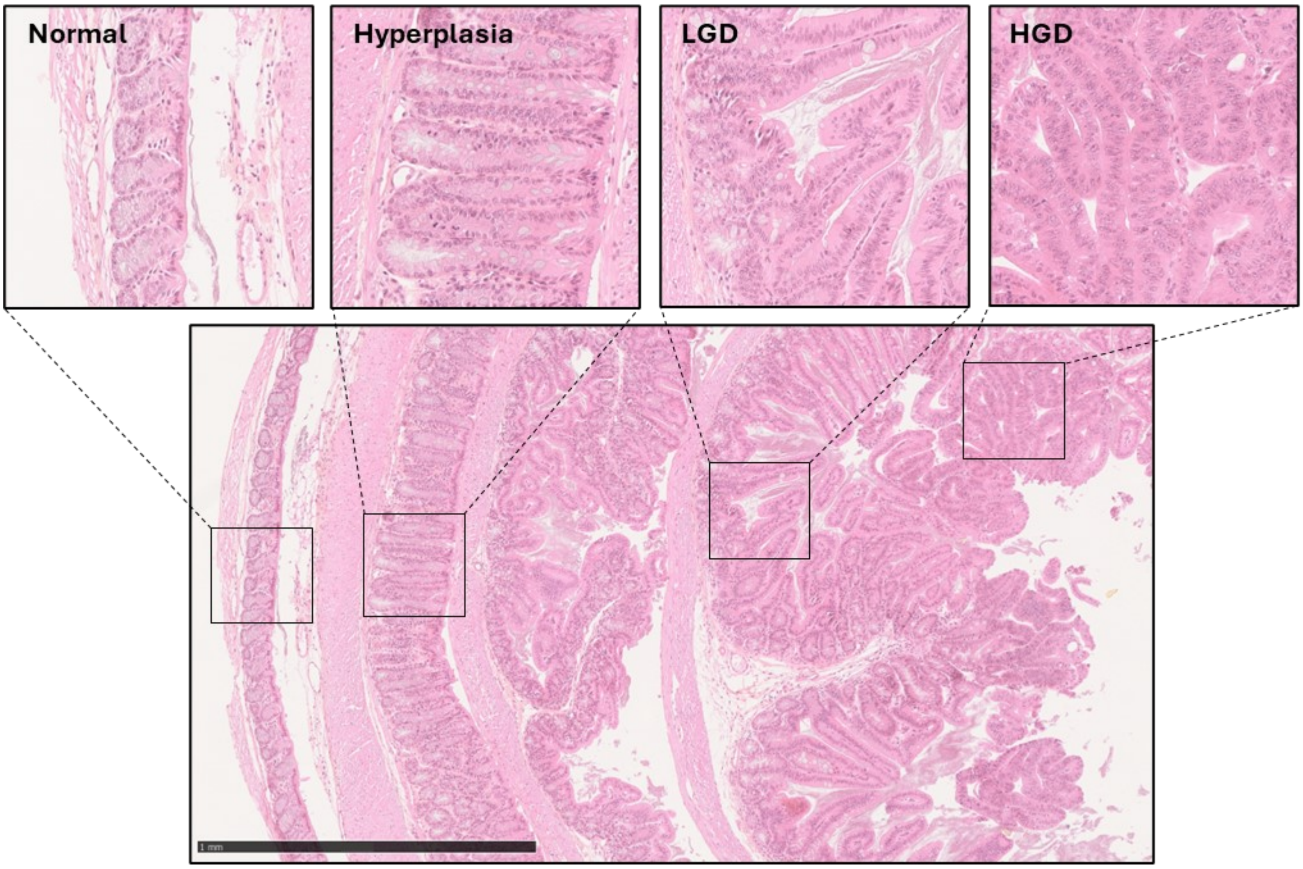

**Figure S1: Representative histopathologic lesions observed in KPC:APC mouse colons.**

Examples of the lesions quantified in the study, including normal epithelium, hyperplasia, low-grade dysplasia (LGD) and high-grade dysplasia (HGD). Scale bar = 1mm.

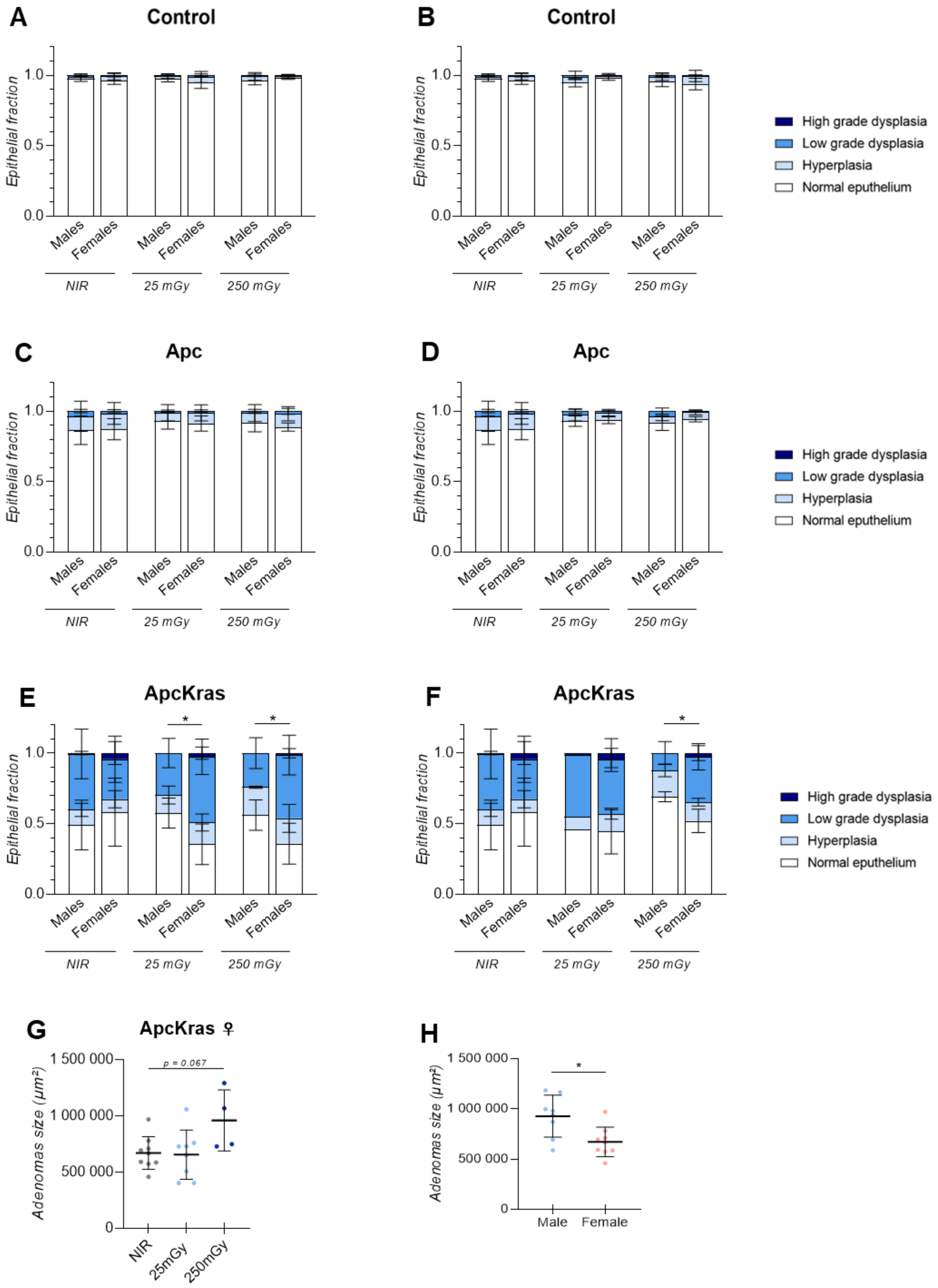

**Figure S2: Sex-specific differences in lesion distribution and adenoma size in KPC:APC mice.**

A–F: Sex-dependent distribution of pre-cancerous lesions in Control (A, B), Apc (C, D), and ApcKras (E, F) mice following early (A, C, E) or late (B, D, F) irradiation. G: Adenoma size in female ApcKras mice after early irradiation. H: Comparison of adenoma size between male and female ApcKras mice under non-irradiated (NIR) conditions.

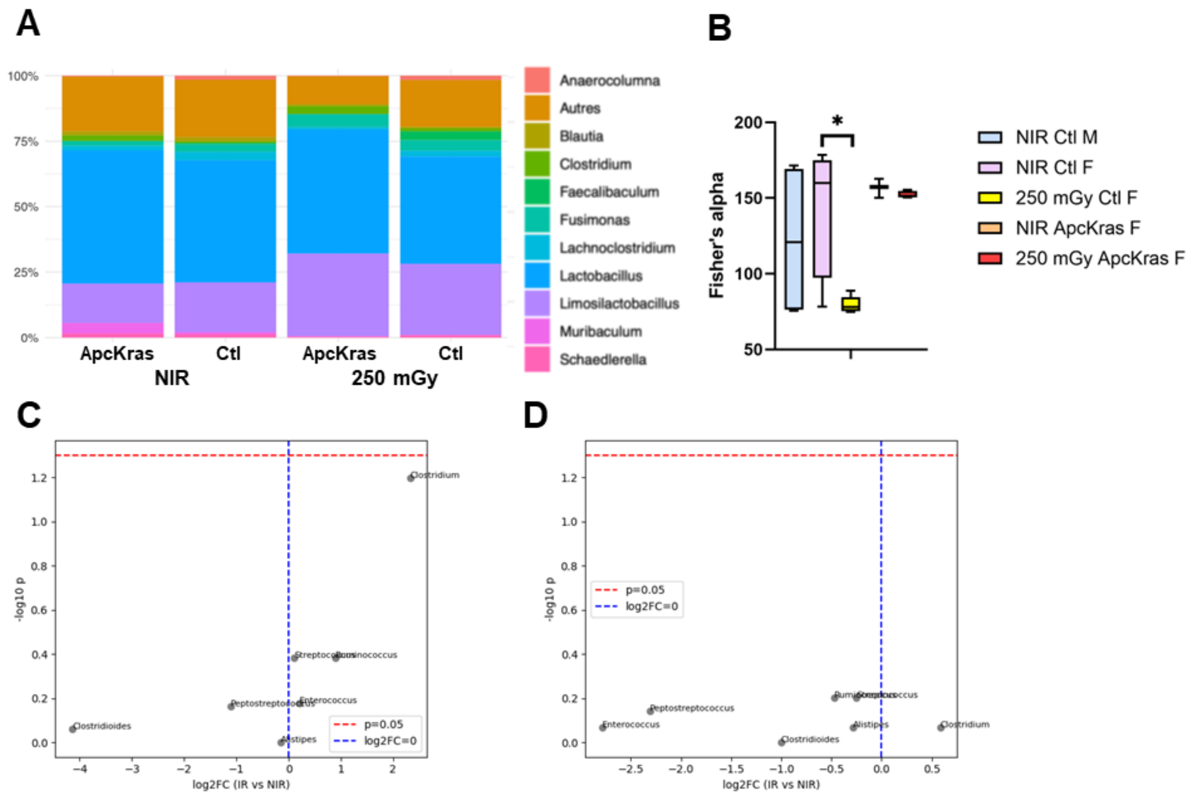

**Figure S3: Gut microbiota modifications after radiation exposure.**

A-B: Distribution of bacterial genera (A) and Fisher's alpha diversity (B) in feces obtained from non-irradiated (NIR) and irradiated (250 mGy) mouse colons. D-E: Radiation-induced modulation of cancer-panel genera after 250 mGy irradiation in [Control] (C) and [ApcKras] animals (red dashed line: p=0.05, blue dashed line: $log_2$ FC = 0).

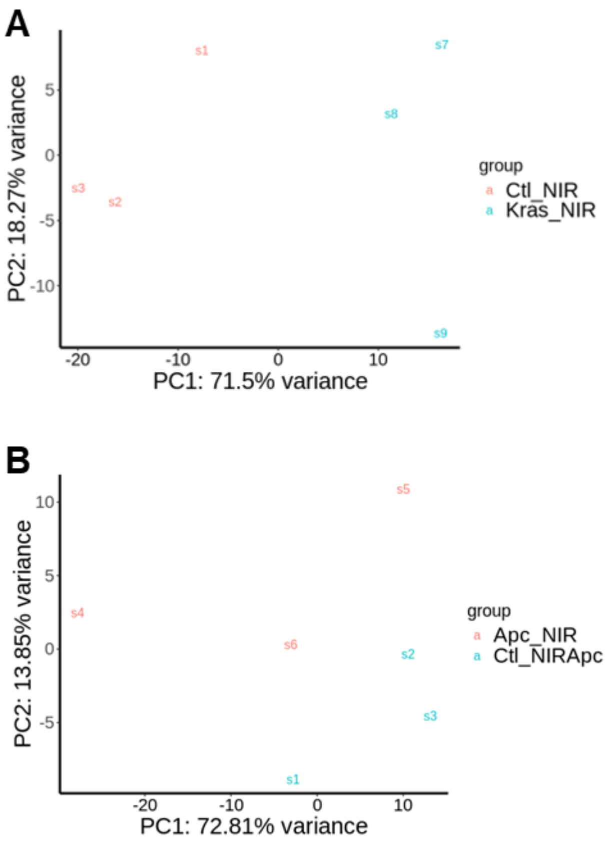

**Figure S4: Principle component analysis (PCA) for the top 1,000 bins ranked by count number after meRIP-seq.**

Principle component analysis (PCA) for the top 1,000 bins ranked by count number, comparing [Control] mice with [ApcKras] (A) or [Apc] (B) non-irradiated mice.

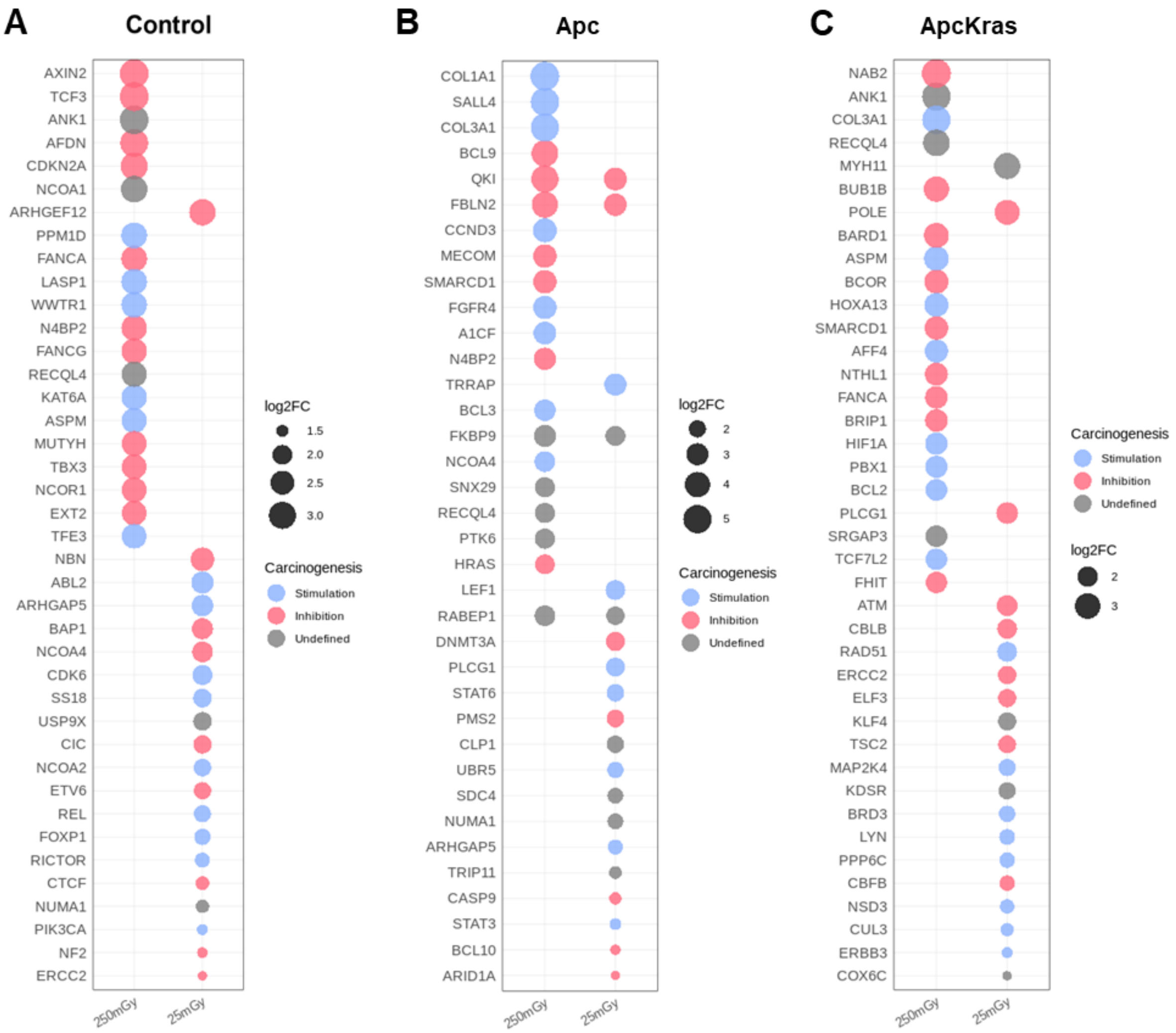

**Figure S5: Modulation of cancer-associated proteins after LMDR in KPC:APC organoids.**

Top 20 dysregulated proteins per irradiation dose encoded by oncogenes and tumor suppressors in [Control] (A), [Apc] (B) and [ApcKras] (C) organoids.

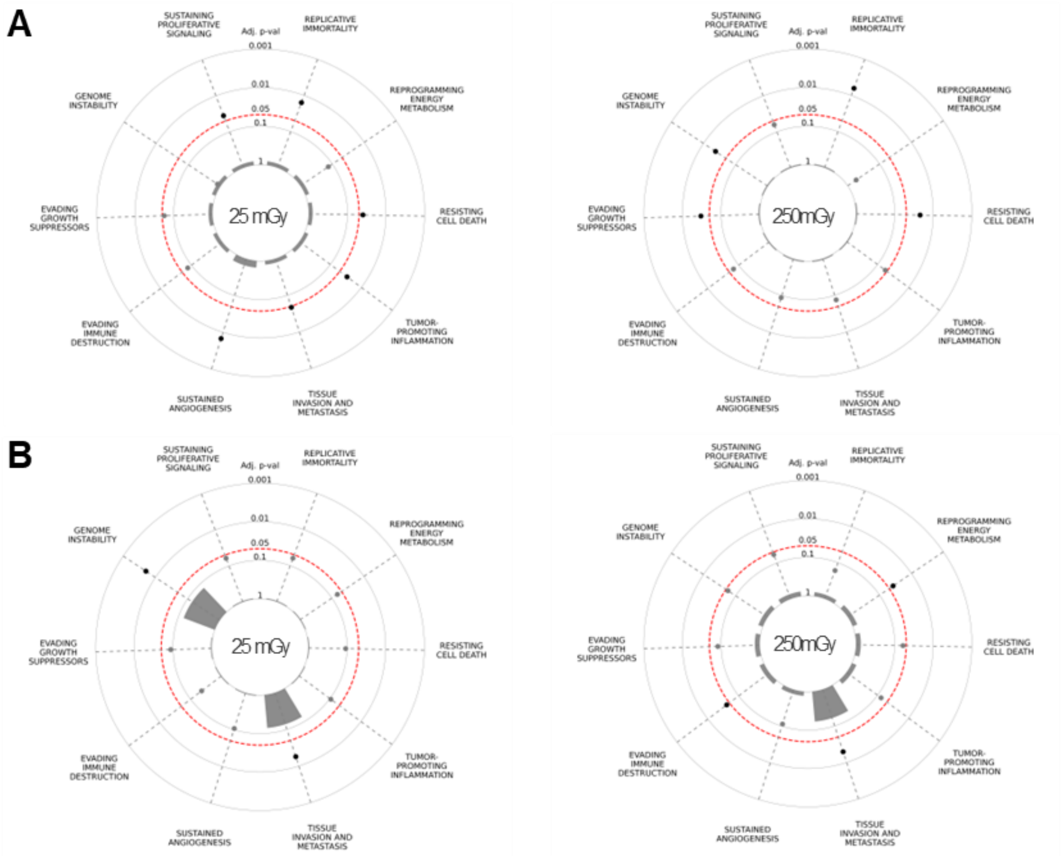

**Figure S6: Overrepresented cancer hallmarks in [Control] and [Apc] organoids**

Overrepresentation of cancer hallmark genes encoding differentially expressed proteins in [Control] (A) and [Apc] (B) organoids following LMDR exposure.

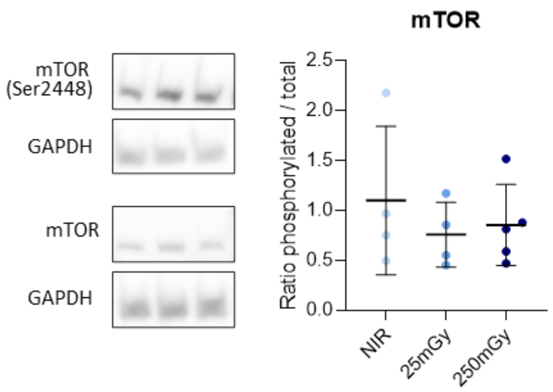

**Figure S7: mTOR phosphorylation in [ApcKras] organoids**

Evaluation of mTORC1 phosphorylation by western blot analysis in [ApcKras] organoids.

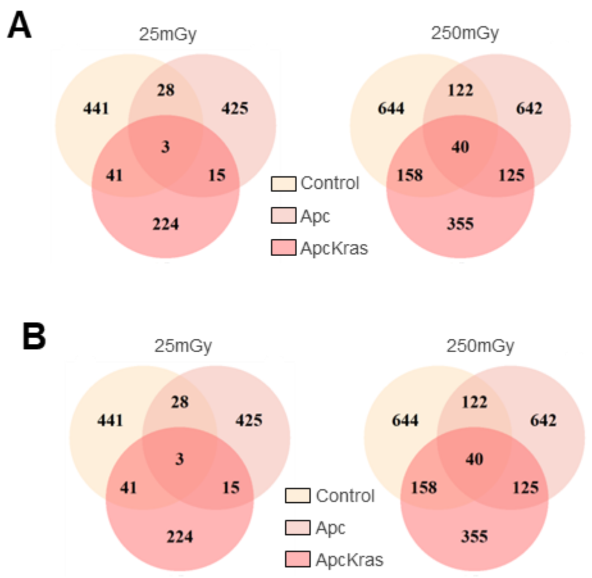

**Figure S8: Overlap of gene and protein regulations after LMDR exposure**

Venn diagrams showing the overlap of DE genes (A) and proteins (B) between [Control], [Apc] and [ApcKras] organoids after LMDR exposures.

#### 2. Supplementary tables

| Sustaining proliferative signaling | Genome instability | Evading growth suppressors | Evading immune destruction | Sustained angiogenesis | Tissue invasion and metastasis | Tumor-promoting inflammation | Resisting cell death | Reprogramming energy metabolism | Replicative immortality |
|---|---|---|---|---|---|---|---|---|---|
| 6/14 | 1/14 | 5/14 | 1/14 | 4/14 | 5/14 | 3/14 | 5/14 | 3/14 | 1/14 |
| RAP2A<br>RAB5C<br>GNA14<br>HSPB1<br>SDC1<br>E2F5 | EXD2 | RAP2A<br>RBM38<br>RAB5C<br>GNA14<br>E2F5 | GNA14 | GNA14<br>SDC1<br>HSPB1<br>EDA2R | RAB5C<br>GNA14<br>HSPB1<br>SDC1<br>E2F5 | RAB5C<br>GNA14<br>SDC1 | HSPB1<br>BOK<br>EDA2R<br>SDC1<br>E2F5 | ACSS2<br>COX7A2L<br>BPGM | E2F5 |

**Table S1: Cancer hallmarks associated with demethylated transcripts in [ApcKras] organoids after 250 mGy irradiation.**

| | **CDS** | **5’ UTR** | **3’ UTR** | **ncRNA** |
|---|---|---|---|---|
| **[Control]** | 33% | 3% | 29% | 35% |
| **[Apc]** | 32% | 9% | 19% | 39% |
| **[ApcKras]** | 26% | 4% | 28% | 42% |

**Table S2: Distribution of the DM peaks in each genomic annotation.**

### 3. Supplementary protocol

#### Gut microbiota analysis

Stools from Kras$^{+/f}$ Apc$^{f/f}$ Cre$^{ERT2+}$ ([ApcKras]) and Kras$^{+/f}$ Apc$^{f/f}$ ([Control]) irradiated (250 mGy) and non-irradiated mice were collected and snap-frozen at the same time these mice were prepared for histopathological analysis (8-12 weeks after tamoxifen administration). Total fecal DNA was extracted using the PureLink Microbiome DNA Purification Kit (Invitrogen, Waltham, MA, USA). Metagenomic sequencing was performed by Macrogen Europe (Amsterdam, NL). Briefly, library construction was performed using the Rapid sequencing DNA – 16S Barcoding Kit 24 V14 (SQK-16S114.24, Oxford Nanopore Technologies ONT, Oxford, UK) according to the manufacturer's instructions. The libraries were loaded onto a MinION/GridION Flow Cell (ONT), sequenced using a GridION Nanopore long-read sequencer (ONT) and acquired by MinKNOW software. Taxonomic annotations and analysis were performed with Kraken2 and KrakenTools [1]. A colorectal cancer-associated panel of bacterial genera was curated from the literature, among them seven genera were detected: Enterococcus, Streptococcus, Peptostreptococcus, Alistipes, Ruminococcus, Clostridium, Clostridioides.
